# Microglial metabolic reprogramming drives cognitive decline in heart failure with preserved ejection fraction

**DOI:** 10.1101/2025.09.10.675257

**Authors:** Swapna Patil, Connor Lantz, Angela Marinovic, Ripon Sarkar, Bo Ryung Lee, Matthew DeBerge

## Abstract

Heart failure with preserved ejection fraction (HFpEF) is a rapidly growing public health concern and an emerging contributor to dementia, yet the mechanisms linking cardiometabolic dysfunction to neurodegeneration remain poorly understood. Here, we demonstrate that HFpEF drives a sustained neuroinflammatory state through microglial metabolic reprogramming. Using a clinically relevant murine model of HFpEF, we identified robust induction of HIF-1α signaling in microglia via integrated transcriptomics and metabolomics, coupled with increased glycolytic metabolism revealed by extracellular flux analysis. Conditional deletion of *Hif1a* in microglia during HFpEF attenuated neuroinflammation, preserved white matter integrity, and rescued cognitive performance. We further identify Sema4D as a HIF-1α–dependent, microglia-derived effector linking metabolic stress to white matter injury. These findings establish a mechanistic bridge between cardiovascular disease and cognitive dysfunction and reveal microglial HIF-1α signaling as a tractable therapeutic strategy for preventing cognitive decline in cardiometabolic disease.

## Introduction

Vascular dementia (VaD) is the second most common form of dementia and frequently co-occurs with other neurodegenerative conditions such as Alzheimer’s disease (AD) (1). While AD is driven largely by intrinsic neuropathological processes, VaD is more closely associated with systemic cardiometabolic dysfunction, where brain injury is thought to result from chronic exposure to extracerebral insults (2, 3). Among the systemic drivers of cognitive decline, heart failure with preserved ejection fraction (HFpEF) has emerged as a particularly concerning contributor.

HFpEF is the most prevalent form of heart failure and is rising in incidence alongside obesity, hypertension, and metabolic syndrome (4, 5). Despite its increasing burden, HFpEF lacks effective treatments and is frequently observed in patients with VaD, where it is associated with the highest mortality among cognitively impaired individuals (6). Although HFpEF and dementia share common vascular risk factors such as aging, hypertension, and hyperlipidemia (7, 8), emerging evidence suggests that HFpEF independently contributes to dementia risk. Compared to patients with only dementia, dementia patients with HFpEF show distinct patterns of brain atrophy and white matter injury, along with worsened depression and reduced quality of life (9-11), effects observed even after adjusting for traditional risk factors. Notably, cognitive decline and gray matter loss in HFpEF occur without changes in cerebral blood flow (12), suggesting mechanisms beyond hypoperfusion. However, the cellular and molecular pathways linking systemic cardiometabolic stress to brain dysfunction remain poorly defined.

Microglia, the resident immune cells of the central nervous system (CNS), are highly sensitive to changes in systemic inflammatory and metabolic states (13-15). In neurodegenerative diseases such as AD and multiple sclerosis, microglia undergo metabolic reprogramming that shifts their functional state and contributes to disease progression (16, 17). However, whether and how HFpEF impacts microglial metabolism and function remains unknown. Using a well-established murine model of HFpEF induced by combined hypertension and metabolic stress (18), we identified robust activation of hypoxia-inducible factor 1α (HIF-1α) signaling as a central regulator of microglial metabolic reprogramming. Selective deletion of microglial HIF-1α signaling revealed its necessity for white matter injury and neuroinflammatory changes during HFpEF. As discussed below, these findings uncover a novel mechanism linking cardiometabolic dysfunction to CNS pathology and highlight microglial metabolic adaptation as a potential therapeutic target in HFpEF.

## Results

To model the multiple comorbidities of human HFpEF, including metabolic stress and hypertension, 8 week old mice were treated with a combination of high fat diet and L-NAME, for 5 weeks (18, 19). Control mice were maintained on standard chow and water. Given the link between HFpEF and white matter injury (10, 20), we assessed changes in the corpus callosum, a major white matter tract in the mouse brain. Compared to chow controls, HFpEF mice exhibited increased microglial abundance and reactivity (Figure 1A), along with loss of oligodendrocytes, an indicator of white matter injury (Figure 1B). Because oligodendrocyte loss and white matter injury are associated with cognitive decline (21), we evaluated behavioral performance. HFpEF mice showed elevated anxiety-like behavior in the open field test (Figures 1C, 1D) and impaired memory in the novel object recognition test (Figures 1E, 1F), with consistent effects observed in both female and male mice. Together, these findings show that HFpEF mice recapitulate key features of human VaD, including neuroinflammation, white matter injury, and cognitive decline.

**Figure 1.**
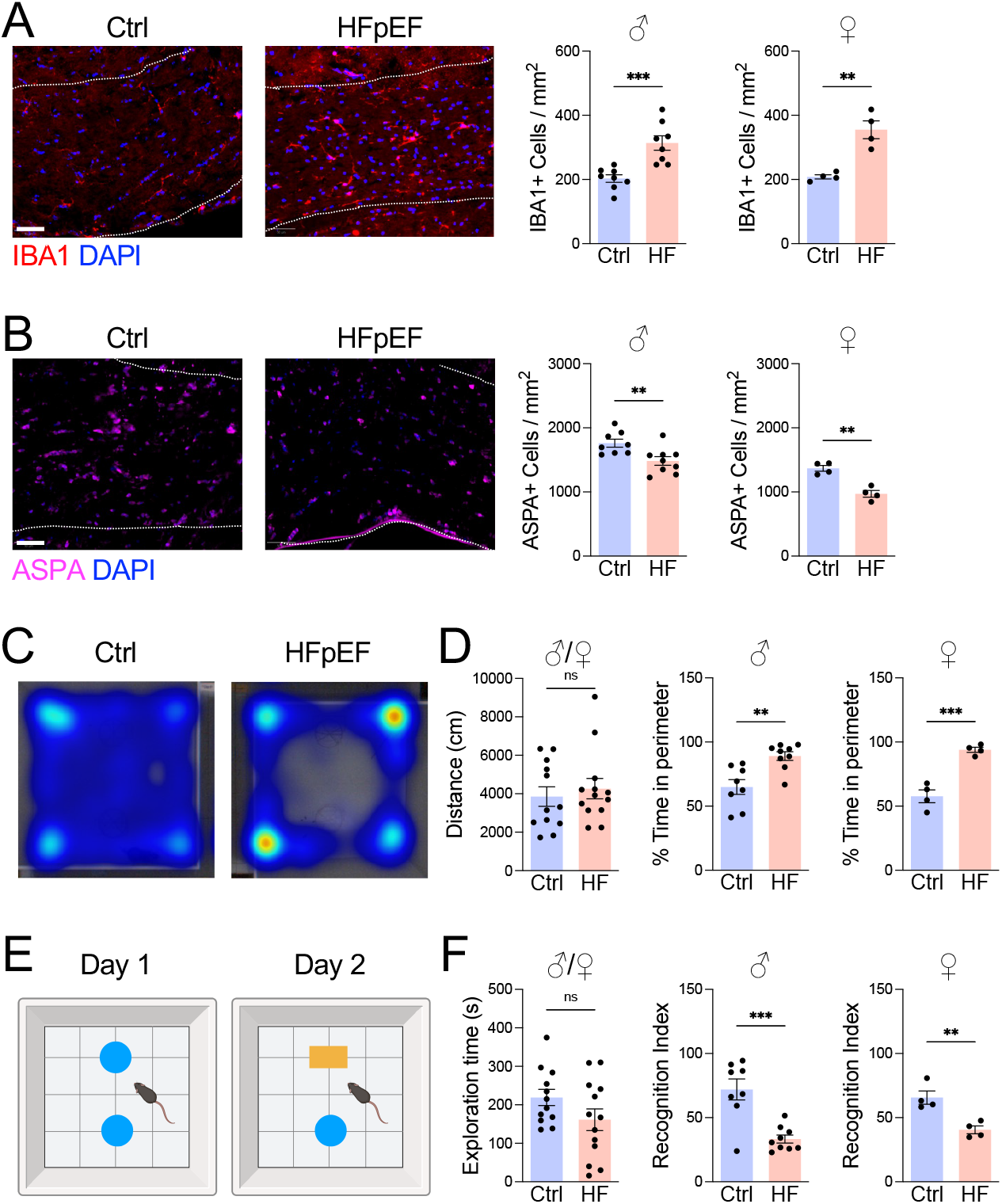
HFpEF causes microgliosis, oligodendrocyte loss, and cognitive decline. **A** Male or female 8-week-old C57BL/6 mice were treated with chow (Ctrl) or HFD+L-NAME (HFpEF) for 5 weeks. **A** IBA1+ microglia and **B** ASPA+ oligodendrocytes measured in corpus callosum (dashed lines). Scale bar, 50 μm. **C** Movement heatmap in open field. **D** Distance traveled and percent time in perimeter of the open field. **E** Schematic of novel object recognition test. **F** Exploration time and percent time with novel object in novel object recognition test. *n =* 4-9 mice/group, *\*P*<0.05, *\*\*P*<0.01, \*\*\**P*<0.001 by unpaired *t* test.

Given the increased abundance of microglia during HFpEF, we next investigated microglia-specific functional changes that may contribute to white matter injury. To do this, we flow sorted microglia from whole cortical tissue after 5 weeks of HFpEF treatment and performed bulk RNA sequencing to assess transcriptional changes (Figure 2A). Microglia were sorted using specific markers, P2RY12 and Tmem119 (22, 23), to exclude CD206^+^ perivascular macrophages and other myeloid populations. Principal component (PC) and heat map analyses revealed clear separation between HFpEF and control groups, indicating distinct transcriptional profiles (Figures 2B, 2C). HFpEF microglia increased expression of inflammatory and glycolytic genes and decreased expression of homeostatic markers (Figures 2D, 2E), consistent with microglial reactivity during HFpEF. Pathway analyses further supported enrichment of inflammatory-related pathways in HFpEF microglia and revealed enrichment of multiple metabolic pathways (Figure 2F), suggesting that HFpEF leads to inflammatory metabolic reprogramming of microglia.

**Figure 2.**
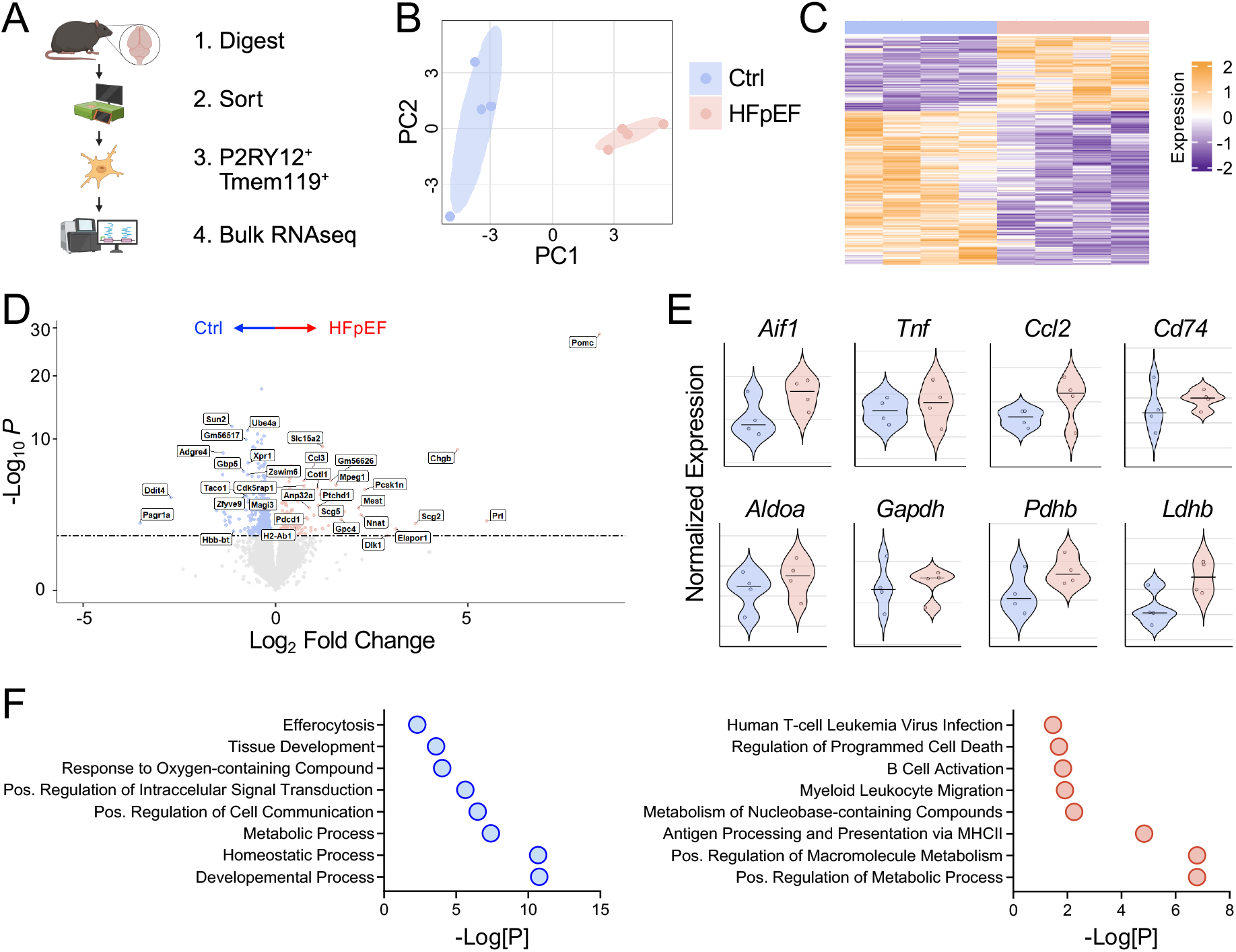
HFpEF induces global transcriptomic reprogramming of microglia. **A** Schematic of microglia transcriptomics after 5 weeks of chow (Ctrl) or HFD+L-NAME (HFpEF) treatment. **B** Principal component (PC) analyses of control and HFpEF microglia. **C** Heatmap of differentially expressed genes (DEG) in control (blue) or HFpEF (red) microglia. **D** Volcano plot of DEGs in microglia. **E** Violin plot of glycolytic and inflammatory genes. **F** Pathways enriched in control (blue) or HFpEF (red) microglia. *n =* 4 mice/group.

To directly measure microglial metabolic reprogramming during HFpEF, we performed paired metabolomic profiling on sorted microglia (Figure 3A). Using untargeted metabolomics, we quantified key intermediates involved in glycolysis, tricarboxylic acid (TCA) cycle, and oxidative phosphorylation. Microglia from HFpEF mice exhibited a distinct metabolic profile (Figures 3B, 3C), characterized by increased levels of glucose-6-phosphate, fructose-1,6-bisphosphate, and lactate (Figure 3D), consistent with enhanced glycolysis. In contrast, TCA intermediates, such as citrate and α-ketoglutarate, were reduced (Figure 3C), suggesting impaired mitochondrial metabolism. Integrated transcriptomic and metabolomic analyses further supported this shift, revealing coordinated enrichment of glycolysis and signaling by hypoxia inducible factor (HIF)-1α (Figure 3E), the master transcriptional regulator of glycolysis (24). Glycolytic network mapping revealed distinct remodeling of glycolytic enzymes in HFpEF microglia (Figure 3F), which was corroborated by quantitative PCR of sorted microglia (Figure 3G). Extracellular flux analysis of sorted microglia confirmed increased glycolysis in HFpEF microglia (Figure 3H), demonstrating glycolytic reprogramming of microglia during HFpEF.

**Figure 3.**
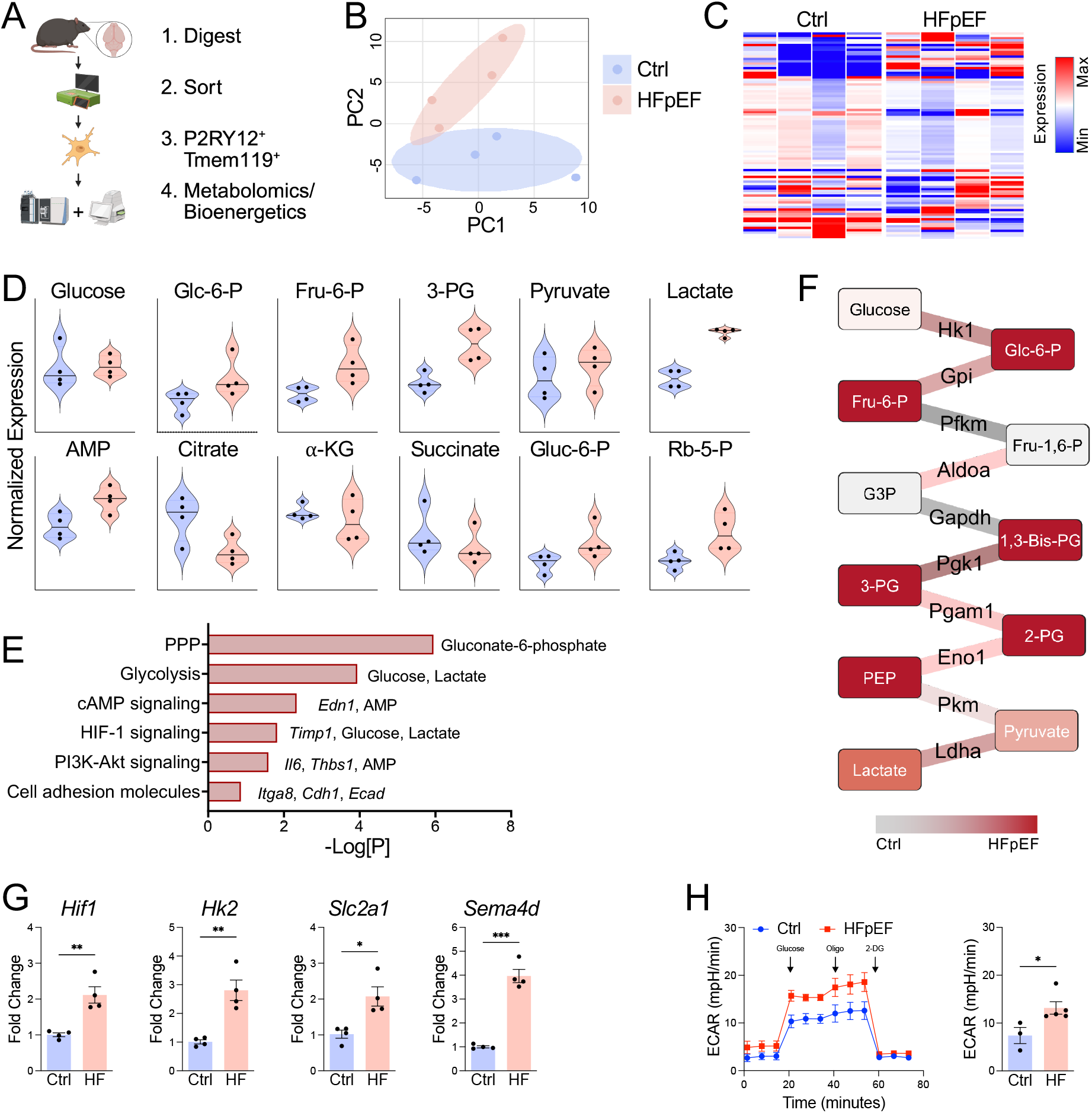
HFpEF drives glycolytic reprogramming of microglia. **A** Schematic of microglia metabolomics after 5 weeks of chow (Ctrl) or HFD+L-NAME (HFpEF) treatment. **B** Principal component (PC) analyses of control and HFpEF microglia. **C** Heatmap of metabolites in control or HFpEF microglia. **D** Violin plot of metabolites. **E** Integrated pathway analysis of upregulated genes and metabolites in HFpEF microglia. **F** Glycolysis network mapping in HFpEF versus control microglia. **G** Gene expression in sorted microglia. **H** Extracellular acidification rate (ECAR) in sorted microglia. *n =* 3-5 mice/group, *\*P*<0.05, *\*\*P*<0.01, \*\*\**P*<0.001 by unpaired *t* test.

Since HIF-1α signaling was enriched in HFpEF microglia, we next directly assessed HIF-1α activation. Using ODD-luciferase HIF reporter mice (Figure 4A), we observed increased bioluminescence signal in the corpus callosum of HFpEF brains compared with controls (Figure 4B), which was accompanied by greater abundance of luciferase-positive microglia (Figure 4C). Because the reporter system does not distinguish between HIF isoforms (25), we performed immunofluorescent staining and observed a marked increase in HIF-1α-positive microglia in HFpEF mice relative to controls (Figures 4D, 4E). Importantly, these findings extended to humans, as cerebellar white matter from HFpEF patients showed a significantly higher abundance of HIF-1α-positive microglia compared with non-HFpEF controls (Figures 4F, 4G). Collectively, these results demonstrate that HFpEF induces robust HIF-1α activation in both mouse and human microglia, suggesting that HIF-1α may serve as a link between metabolic reprogramming and downstream neuropathology.

**Figure 4.**
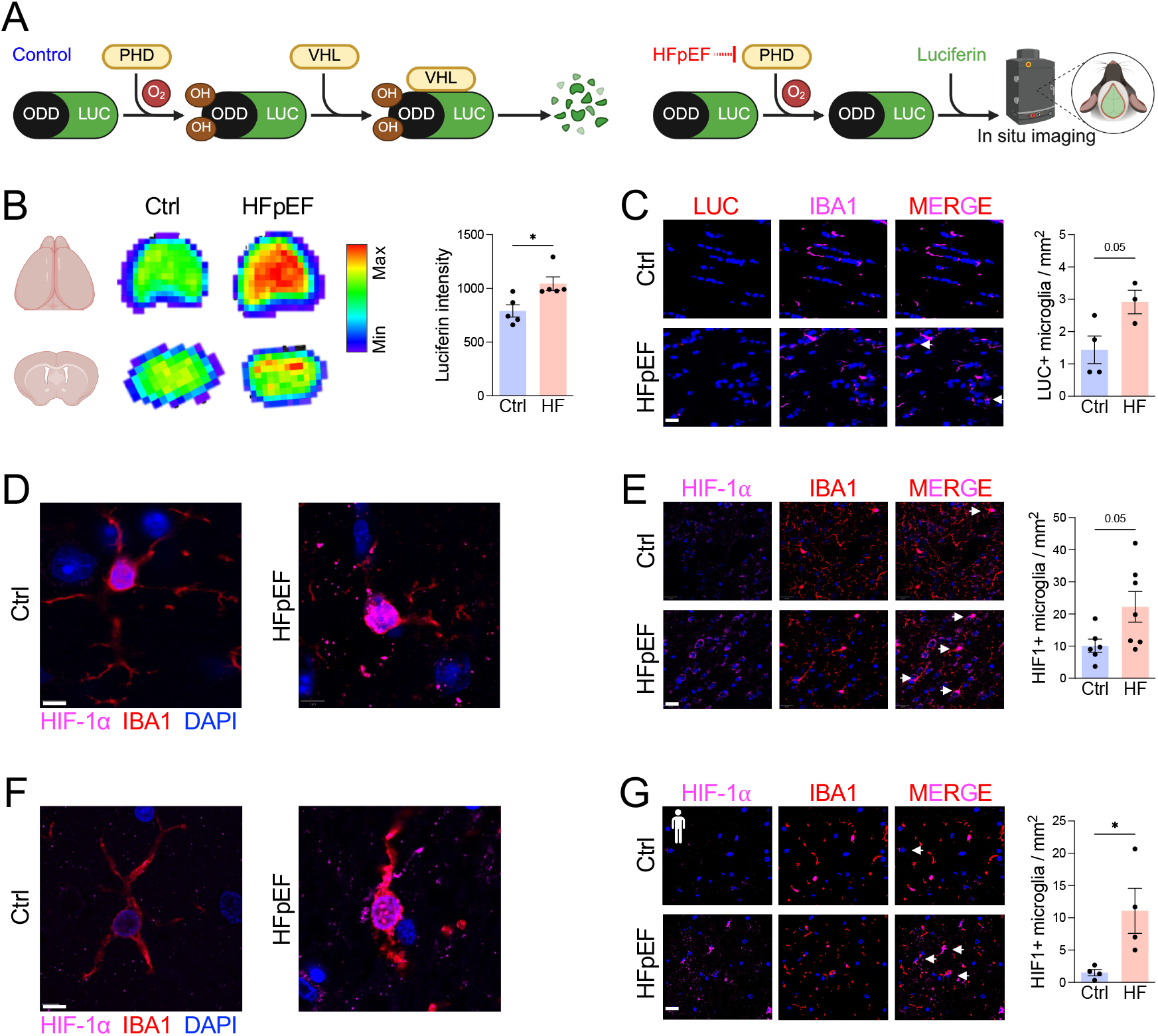
HFpEF induces HIF-1α activation in mouse and human microglia. **A** Schematic of ODD-luciferase HIF reporter mouse. **B** Luciferin signal intensity in control or HFpEF brains. **C** Quantification of luciferase positive microglia. Scale bar, 20 μm. **D** HIF-1α nuclear localization in mouse microglia. Scale bar, 5 μm. **E** Quantification of HIF-1α positive mouse microglia. Scale bar, 20 μm. **F** HIF-1α nuclear localization in human microglia. Scale bar, 5 μm. **G** Quantification of human HIF-1α positive microglia in cerebellar white matter. Scale bar, 20 μm. For **B-E**, *n =* 3-7 mice/group, *\*P*<0.05, *\*\*P*<0.01, \*\*\**P*<0.001 by unpaired *t* test. For **G**, *n* = 4 autopsy cases/group, *\*P*<0.05 by unpaired *t* test.

To determine whether HIF-1α activation in microglia contributed to neuropathology during HFpEF, we generated mice with microglia-specific deletion of *Hif1* (Figure 5A), with knockdown efficiency confirmed in sorted microglia (Figure 5B). Microglial *Hif1* deletion did not alter body weight or blood pressure during HFpEF, indicating no effect on systemic disease severity (Figures 5C, 5D). In contrast, microglial *Hif1*-deficiency preserved oligodendrocyte abundance in the corpus callosum (Figure 5E). While microglia abundance was unchanged (Figure 5E), cortical extracts from microglial *Hif1*-deficient mice exhibited reduced expression of inflammatory genes (Figure 5F), consistent with an inflammatory role for microglia HIF-1α signaling in HFpEF. Functionally, microglial *Hif1*-deficiency rescued HFpEF-associated behavioral deficits, as mice displayed increased locomotor activity and normalized exploratory patterns in the open field test (Figure 5G), along with improved recognition memory in the novel object recognition test (Figure 5H). These findings identify microglial HIF-1α activation as a key step driving white matter injury and cognitive decline in HFpEF.

**Figure 5.**
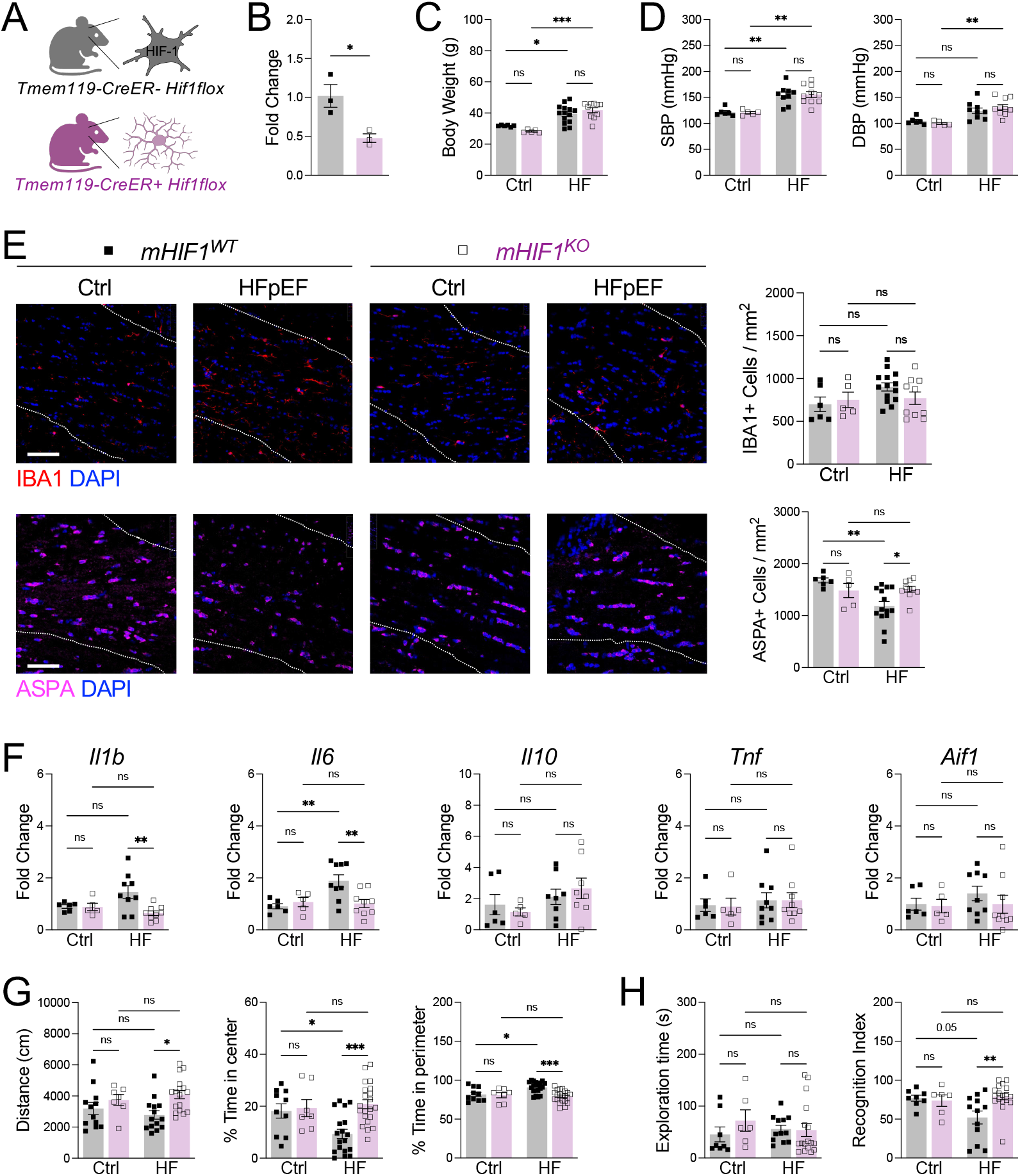
Microglial HIF-1α activation drives neuropathology and cognitive dysfunction during HFpEF. **A** Schematic of microglial *Hif1* deletion. **B** Knockdown of *Hif1* in sorted microglia. **C** Changes in body weight (g). **D** Systolic (SBP) and diastolic (DBP) blood pressure measurements. **E** IBA1+ microglia and ASPA+ oligodendrocytes measured in corpus callosum (dashed lines). Scale bar, 50 μm. **F** Gene expression in cortical extracts. **G** Distance traveled and percent time in center or perimeter of the open field. **H** Exploration time and percent time with novel object in novel object recognition test. For **B**, *n =* 3 mice/group, *\*P*<0.05 by unpaired *t* test. For **C-H**, *n* = 5-18 mice/group, *\*P*<0.05, *\*\*P*<0.01, \*\*\**P*<0.001 by two-way ANOVA followed by Tukey’s test.

Next, we sought to identify the link between microglial HIF-1α activation and white matter injury. A recent genome-wide association study identified *SEMA4D* as a gene linked with all-cause dementia in humans (26). *SEMA4D* encodes semaphorin 4D (Sema4D), a protein elevated in the circulation of HFpEF patients that can induce apoptosis in mature oligodendrocytes (27-29), suggesting it may mediate white matter injury downstream of microglial HIF-1α signaling. Consistent with this hypothesis, sorted microglia from HFpEF mice expressed higher levels of *Sema4d* compared to controls (Figure 3G). To model microglia responses, we used bone marrow-derived macrophages (BMDMs) and treated them with HFpEF-relevant stimuli, including TLR4 agonists or hypoxia (30). Both stimuli robustly induced *Sema4d* expression (Figure 6A), with concomitant increases in Sema4D protein production and secretion (Figures 6B, 6C). *Hif1*-deficient BMDMs exhibited markedly reduced Sema4D induction at both transcript and protein levels (Figures 6D-F), demonstrating that HIF-1α is required for Sema4D expression in myeloid cells. Furthermore, microglial *Hif1*-deficiency blunted HFpEF-induced *Sema4d* expression (Figure 6G), supporting the requirements for HIF-1α in driving microglial *Sema4D* expression.

**Figure 6.**
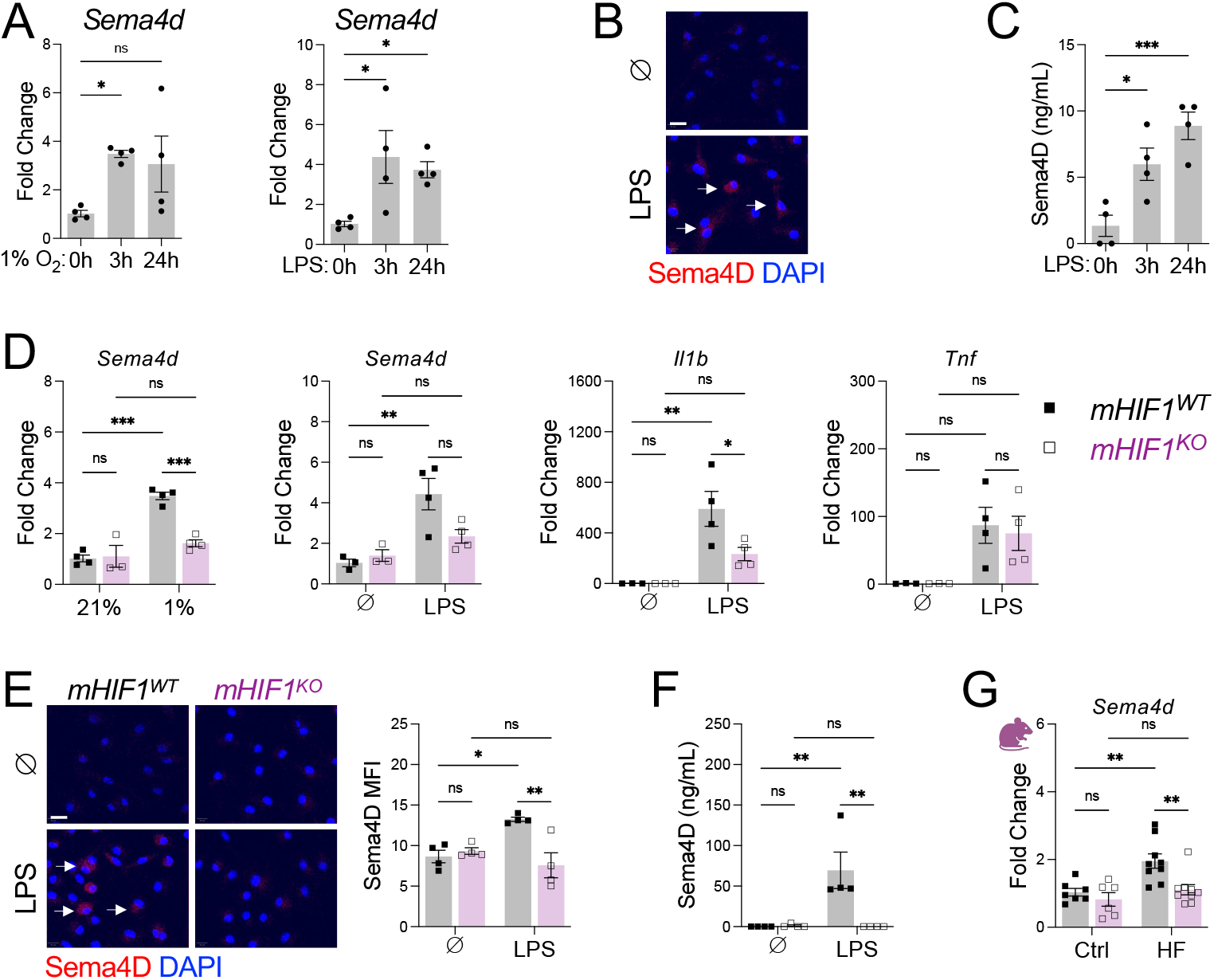
HIF-1α activation is required for microglial Sema4D production. **A** Gene levels of *Sema4d* after hypoxia or LPS treatment in bone marrow derived macrophages (BMDM). **B** Immunofluorescent staining of Sema4D in BMDM after LPS treatment. **C** Soluble Sema4D production by BMDM after LPS treatment. **D** Gene levels in *Hif1*-deficient BMDMs after 3 hours of treatment. **E** Immunofluorescent staining of Sema4D in *Hif1*-deficient BMDMs after 3 hours LPS treatment. **F** Soluble Sema4D production by *Hif1*-deficient BMDMs after 3 hours LPS treatment. **G** Gene levels of *Sema4d* in cortical extracts. For **A-F**, *n* = 3-4 wells/group, *\*P*<0.05, *\*\*P*<0.01, \*\*\**P*<0.001 by one-way or two-way ANOVA followed by Tukey’s test. For **G**, *n =* 5-9 mice/group, *\*\*P*<0.01, \*\*\**P*<0.001 two-way ANOVA followed by Tukey’s test.

Finally, to test whether microglial-derived Sema4D contributes directly to white matter injury and cognitive decline in HFpEF, we generated mice with microglia-specific deletion of *Sema4d* (Figure 7A) with knockdown efficiency confirmed in sorted microglia (Figure 7B). Microglial *Sema4d*-deficiency had no effect on body weight or systolic blood pressure during HFpEF (Figure 7C, 7D), indicating that systemic disease severity was unchanged. In contrast, loss of microglial *Sema4d* preserved oligodendrocyte abundance in the corpus callosum without affecting microglial abundance (Figure 7E). This structural protection translated to improved behavioral outcomes as mice with microglial *Sema4d*-deficiency exhibited normalized exploratory behavior in the open field test (Figure 7F) and improved recognition memory in the novel object recognition test (Figure 7G). Taken together, our findings reveal microglial-derived Sema4D as a central mediator of oligodendrocyte loss and cognitive dysfunction in HFpEF, mechanistically linking HIF-1α-dependent microglial metabolic reprogramming to white matter injury and dementia-related outcomes.

**Figure 7.**
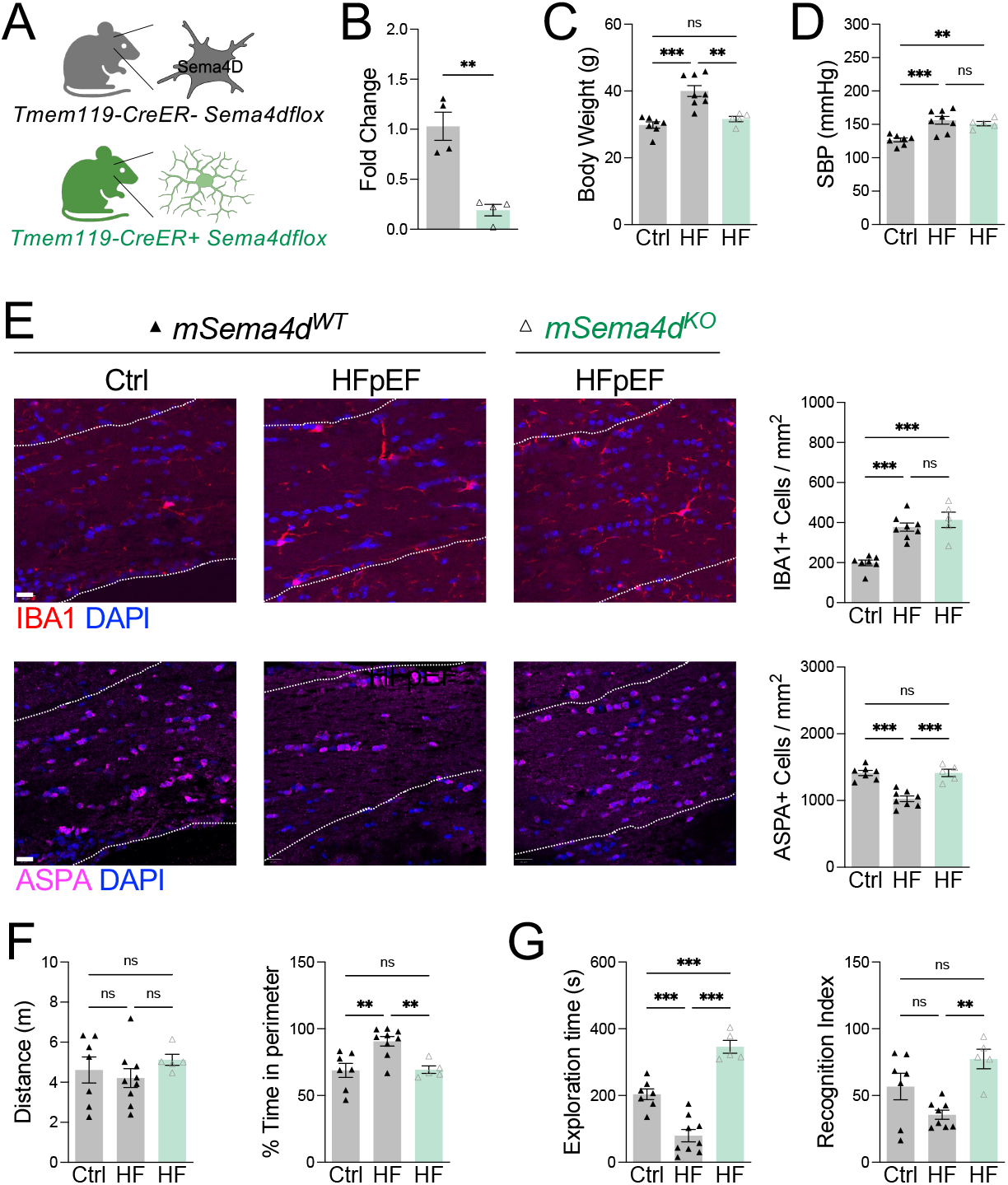
Microglia Sema4D signaling drives oligodendrocyte loss and cognitive impairment during HFpEF. **A** Schematic of microglial *Sema4d* deletion. **B** Knockdown of *Sema4d* in sorted microglia. **C** Changes in body weight (g). **D** Systolic blood pressure (SBP) measurements. **E** IBA1+ microglia and ASPA+ oligodendrocytes measured in corpus callosum (dashed lines). Scale bar, 20 μm. **F** Distance traveled and percent time in perimeter of the open field. **G** Exploration time and percent time with novel object in novel object recognition test. *n =* 5-8 mice/group, *\*P*<0.05 by one-way ANOVA followed by Tukey’s test.

## Discussion

Our data establish a mechanistic pathway by which cardiometabolic stress promotes cognitive impairment through microglial metabolic reprogramming. We show that HFpEF induces a glycolytic shift in microglia, driven by HIF-1α, which was observed in both mouse and human microglia. Microglial *Hif1* deletion reduced neuroinflammation, preserved oligodendrocytes, and rescued behavioral deficits, establishing microglial HIF-1α as a key regulator of neuropathology in HFpEF. We further demonstrate that HIF-1α drives expression of Sema4D and that microglial *Sema4d* deletion protects against oligodendrocyte loss and behavioral impairment. Together, these findings define a microglial HIF-1α-Sema4D axis linking cardiometabolic disease to dementia and support this pathway as a potential therapeutic target for neuroprotection.

Cellular and tissue-level metabolic remodeling has emerged as a central pathological mechanism in HFpEF (31). In the present study, we identified a glycolytic metabolic reprogramming shift in microglia driven by HIF-1α activation. HFpEF microglia showed a coordinated upregulation of glycolytic genes, including key enzymes such as *Slc2a1, Hk2, Pkm*, and *Ldhb*, and accumulation of glycolytic intermediates, including glucose-6-phosphate, fructose-1,6-bisphosphate, and lactate, accompanied by a reduction in TCA cycle metabolites such as citrate and α-ketoglutarate, indicating impaired oxidative phosphorylation. Extracellular flux analysis confirmed increased glycolytic flux, establishing HIF-1α as a central regulator of this proinflammatory metabolic shift. By contrast, cardiac macrophages in HFpEF preferentially engage fatty acid oxidation and mitochondrial respiration to support tissue remodeling and hematopoietic stem cell activation (19, 32), highlighting a striking divergence in metabolic adaptations among myeloid populations across tissues. These findings reveal that systemic cardiometabolic stress drives tissue-specific metabolic programs with HIF-1α-dependent glycolysis in microglia contributing to neuroinflammation, white matter injury, and cognitive impairment.

To identify downstream effectors of microglial metabolic reprogramming, we focused on Sema4D, recently identified as a novel locus for all-cause dementia (26). HFpEF microglia showed robust, HIF-1α-dependent upregulation of *Sema4d*, consistent with its regulation by HIF-1α in the tumor microenvironment (33), and linking metabolic reprogramming to effector function. Strikingly, microglia-specific *Sema4d* deletion preserved oligodendrocytes without altering microglia abundance, supporting a model in which microglial-derived Sema4D acts directly on oligodendrocytes to promote white matter injury. This is further supported by our recent work showing enhanced microglia-oligodendrocyte crosstalk via Sema4D following myocardial infarction (34), indicating a conserved injury response that may link cardiovascular stress to neurodegeneration. Notably, Sema4D is also increased in the brain during other neurodegenerative processes, including Huntington’s Disease and AD (35), raising the possibility that cardiometabolic stress interacts with established drivers of neurodegeneration, such as tau and amyloid-β, to shape disease outcomes.

In a therapeutic context, our findings suggest that clinical inhibition of the microglial HIF-1α-Sema4D axis may reduce white matter injury and cognitive decline during cardiometabolic disease. Highly selective HIF-1α inhibitors have demonstrated favorable safety profiles and sustained activity in clinical trials for the treatment of cancer in humans (36). Since HIF-1α exerts a protective role in other CNS populations (37), targeted delivery to microglia will be essential to maximize efficacy and minimize off-target effects (38). In contrast, Sema4D inhibition may yield more immediate clinical benefit. Pepinemab, an antibody targeting Sema4D, is currently in clinical trials for Huntington’s Disease and AD (39, 40). In mouse models of neurodegeneration, it has been shown to reduce neuroinflammation, stabilize the blood-brain barrier, and protect oligodendrocytes (28), suggesting broader neurovascular benefits. Because most dementias are driven by modifiable cardiometabolic risk factors (41), future studies should also investigate how lifestyle interventions, as well as current HFpEF therapies such as SGLT2 inhibitors (42), influence microglial metabolic reprogramming and therapeutic outcomes.

A limitation of our study is that experiments were performed in relatively young adult mice, and age-related changes could modulate microglial responses and the benefits of targeting microglial metabolic reprogramming (43). Studying younger animals allowed us to define how cardiometabolic stress disrupts a healthy neuroimmune environment, which remains clinically relevant given the links between cardiometabolic disease and white matter injury in adolescent and adult humans (44, 45). Similarly, because Sema4D can signal through multiple cell types (46), cell-specific receptor knockouts will be critical to clarify contributions of oligodendrocyte signaling versus other CNS populations. Finally, evaluating therapeutic potential in mice with established disease using temporal deletion or pharmacologic inhibition will help model clinical interventions more accurately.

In conclusion, our findings reveal that metabolic reprogramming of microglia by HIF-1α links cardiometabolic disease to white matter injury and cognitive decline through Sema4D production. The newly discovered roles for HIF-1α and Sema4D provide a framework for understanding how systemic disease drives neurodegeneration and opens new avenues for therapeutic intervention aimed at targeting microglial metabolism and effector signaling to attenuate white matter injury and preserve cognitive function.

## Methods

### Mice

*C57BL/6* mice were originally purchased from Taconic Biosciences. *Cx3cr1CreER* (B6.129P2(C)-*Cx3cr1*^*tm2*.*1(cre/ERT2)Jung*^/J, stock no: 020940), *Tmem119-2A-CreERT2* (C57BL/6-*Tmem119*^*em1(cre/ERT2)Gfng*^/J, stock no: 031820), and *Hif1a* flox (B6.129-*Hif1a*^*tm3Rsjo*^/J, stock no: 007561) were originally purchased from Jackson Laboratories. *Sema4d* flox (Cat no. NM-CKO-2116132) were purchased from Shanghai Model Organisms. HIF reporter (ODD-luciferase) mice were generously provided by Holger Eltzschig (UTHealth-Houston) (25, 47). All mice were bred in our animal facility before use. To generate mice with microglia-specific gene deletion, flox mice were bred with *Cx3cr1CreER* or *Tmem119-2A-CreERT2* mice. To induce Cre-recombinase activity, mice received tamoxifen (i.p. 75 mg/kg per mouse) dissolved in corn oil for 5 consecutive days (48). After the final injection, mice were rested for 30 days to ensure turnover of monocytes and macrophages (49). For controls, Cre-negative littermates were cohoused with Cre-positive mice and treated with tamoxifen. Mice were housed in a temperature- and humidity-controlled pathogen-free environment and kept on a 12h/12h day/night cycle with an ambient temperature of 22° C. Animal studies were conducted in accordance with guidelines using a protocol approved by the Animal Welfare Committee at UTHealth-Houston. Two- to four-month female and male mice were used for experiments.

### HFpEF Model

To induce HFpEF, mice were treated with high fat diet (HFD, 60% kcal fat) combined with pH adjusted drinking water supplemented with 0.5 grams/liter L-NAME for 5 weeks as previously described (18, 19). HFD and L-NAME drinking water was changed twice per week. Control mice were maintained on standard chow and water. Body weight was recorded weekly throughout the study to track weight gain, and mice that failed to gain weight over the course of the treatment period were excluded from analyses.

### Luciferase Activity Imaging

ODD-luciferase HIF reporter mice were injected intraperitoneally with 100 mg/kg D-luciferin and euthanized 10 minutes later. Brains were rapidly removed without perfusion and imaged using the bioluminescence mode of the IVIS Lumina III system with serial exposure times. Regions of interest were defined and bioluminescence intensity was quantified using Living Image software.

### Primary Microglia Isolation

Primary microglia were isolated from adult (2 to 4 months old) mouse brains after 5 weeks of HFpEF or control treatment. Following euthanasia, mice were perfused with 20 mL of ice-cold PBS to remove peripheral blood and immune cells. Whole brains were rapidly removed, meninges were carefully resected, and tissue was minced and enzymatically digested using the Neural Tissue Dissociation Kit (P) (Miltenyi) by incubating at 37°C on a thermal mixer (Thermo Scientific) set to shake at 300 rpm for 20 minutes. The resulting single cell suspension was passed through a 100 μm cell strainer and resuspended in HBSS with 40% Percoll. Myelin was removed by centrifuging the cell suspension at 700 x g for 25 minutes at 4°C without brake, allowing efficient separation of myelin debris from the cell pellet. Cells were then washed and resuspended in FACS buffer (PBS with 2% FBS and 2 mM EDTA) and blocked with TruStain FcX (1:100 dilution) antibody on ice for 10 minutes. For staining, cells were labeled with Tmem119-Alexa Fluor 488, P2RY12-PE, CD206-APC, Ly6C-APC, Ly6G-APC, and CD11b-APC/Cy7 (all 1:300 dilution) in FACS buffer for 15 minutes in the dark on ice. After sequential washes, cells were resuspended in MACSQuant Tyto Running Buffer (Miltenyi) and viability was assessed by the addition of DAPI (1:1000 dilution). The stained cell suspension was loaded into a MACSQuant Tyto High-Speed Cartridge (Miltenyi) and microglia were isolated by gating on live, CD11b^+^, Tmem119^+^, and P2RY12^+^ cells. Perivascular macrophages and other myeloid cells, including monocytes and neutrophils, were excluded using a dump channel (CD206, Ly6C, Ly6G). Microglia were recovered from the positive collection chamber and used for downstream applications.

### Primary Microglia Culture

Primary microglia were isolated as described above, with cells from approximately three to five mice per group typically pooled for each preparation. After myelin debris removal, microglia were purified using CD11b-positive magnetic selection (BioLegend). Cells were plated in T25 flasks in 10 mL of DMEM supplemented with 10% FBS, 1% penicillin/streptomycin, 1% L-glutamine, and 100 ng/mL M-CSF (50). Fresh medium was added on days 2 and 5, and Cre-mediated recombination was induced beginning on day 7 of culture by adding 1 μM tamoxifen (Sigma) to the culture media. Microglia were maintained until days 10–14, at which point they were harvested and seeded at 5×10^4^ cells per well in 96-well tissue culture–treated plates for experiments.

### Seahorse Analysis

Bioenergetic function was measured in sorted primary microglia by Seahorse extracellular flux analysis. Purified microglia were plated on Cell-Tak–coated XFe96 culture plates (Corning) at a density of 150,000 cells per well and allowed to adhere for 2 hours prior to assay. Measurements were conducted in pre-warmed, phenol red–free DMEM lacking glucose and sodium pyruvate (Millipore Sigma), supplemented with 2 mM L-glutamine, pH 7.4. Baseline readings were collected in triplicate, followed by sequential injection of the following compounds: (1) glucose (25 mM) to stimulate glycolysis; (2) oligomycin (1.5 μM) to inhibit ATP synthase; (3) sodium pyruvate (1 mM) with FCCP (1.5 μM) to uncouple oxidative phosphorylation; and (4) rotenone (100 nM) and antimycin A (1 μM) to block complexes I and III, together with 2-deoxyglucose (500 mM) to inhibit glycolysis. Extracellular acidification rate (ECAR) and oxygen consumption rate (OCR) were recorded using a Seahorse XFe96 Analyzer (Agilent), and Wave software (Agilent) was used to calculate glycolytic flux and mitochondrial respiration.

### RNA Transcriptomics

Bulk RNA sequencing was performed at the Cancer Genomics Center at UTHealth. RNA integrity was assessed using an RNA integrity number (RIN) assay, and only samples with RIN values greater than 7 were processed. Ultra–low input library preparation was performed, and sequencing was carried out on an Illumina NextSeq platform at a depth of ∼20 million paired-end reads (150 bp) per sample. FASTQ files storing the raw sequencing reads were processed using the nf-core/rnaseq NextFlow pipeline (v3.19.0) (51). Briefly, adapter sequences and low-quality bases were trimmed using Trim Galore! (v0.6.10) (52) and sequence quality was assessed with FastQC (v0.12.1). Reads were then aligned to the *Mus musculus* reference genome (GRCm39; Ensembl release 114) using STAR (53) and gene expression was quantified using Salmon (54), generating a raw counts matrix across all samples. The resulting counts matrix was imported into R for downstream analyses. Normalized counts were obtained using the DESeq2 (55) variance-stabilizing transformation (vst) to correct for sequencing depth and biases. Principal component analysis (PCA) was performed on the top 500 most variable genes to assess sample clustering, and visualizations were generated using ggplot2 (v3.5.2). Differential expression analysis was carried out in DESeq2 using the Wald test with Benjamini–Hochberg correction. Genes were considered differentially expressed if they exhibited an adjusted *p*-value (FDR) less than 0.05 and an absolute log2 fold-change greater than 0.5. Pathway enrichment analysis was performed by g:Profiler (v0.2.3) (56) on sets of genes identified between HFpEF and control groups. Genes were stratified by direction of change of the log2 fold-change and an adjusted p-value less than 0.1 to identify pathways enriched in either control or HFpEF groups respectively.

### Metabolomics

To determine the relative abundance of polar metabolites in primary mouse microglia samples, extracts were prepared and analyzed by ultra-high-resolution mass spectrometry (HRMS) (57). Approximately 100,000 microglia were snap frozen in liquid nitrogen, and metabolites were extracted using ice-cold 80/20 (v/v) methanol/water with 0.1% ammonium hydroxide. Extracts were centrifuged at 17,000 g for 5 min at 4°C, and supernatants were transferred to clean tubes, followed by evaporation to dryness under nitrogen. Dried extracts were reconstituted in deionized water, and 10 μL were injected for analysis by Ion chromatography (IC)-HRMS. IC mobile phase A (MPA; weak) was water, and mobile phase B (MPB; strong) was water containing 100 mM KOH. A Thermo Scientific Dionex ICS-6000+ system included a Thermo IonPac AS11 column (4 μm particle size, 250 x 2 mm) with column compartment kept at 35°C. The autosampler tray was chilled to 4°C. The mobile phase flow rate was 360 μL/min, and the gradient elution program was: 0-2 min, 1% MPB; 2-25 min, 1-40% MPB; 25-39 min, 40-100% MPB; 39-50 min, 100% MPB; 50-50.5, 100-1% MPB. The total run time was 55 min. To enhance the desolvation for better sensitivity, methanol was delivered by an external pump and combined with the eluent via a low dead volume mixing tee. Data were acquired using a Thermo Orbitrap IQ-X Tribrid Mass Spectrometer under ESI negative ionization mode.

### Integrated Transcriptomics and Metabolomics Analyses

To integrate transcriptomic and metabolomic profiles, normalized gene expression (variance-stabilized counts) and metabolite abundances (log-transformed, batch-corrected intensities) were aligned by sample identifiers. Concordance between molecular layers was assessed through correlation analyses, where metabolite abundances were correlated with the expression of genes involved in the same pathways (e.g., glycolysis and macrophage inflammatory signaling). Significant gene–metabolite pairs were identified using Pearson correlation coefficients with false discovery rate (FDR) adjustment. Pathway-level integration was performed using MetaboAnalyst 6.0 (Joint Pathway Analysis module) (58). Specifically, differentially expressed genes and differentially abundant metabolites (adjusted p < 0.05) were uploaded, and joint enrichment analysis was carried out using the KEGG Mus musculus pathway library. Gene and metabolite data were weighted equally, and significance of pathway enrichment was assessed using hypergeometric testing with multiple-testing correction. To further explore cross-omic networks, fold changes for both transcripts and metabolites were imported into Cytoscape (v3.10.0) and mapped onto the glycolysis/gluconeogenesis pathway using the KEGG plugin.

### Non-Invasive Blood Pressure

Noninvasive blood pressure was measured in conscious mice using the tail-cuff method with a CODA system (Kent Scientific). To minimize stress-related variability, mice were acclimated to the restrainer and tail-cuff apparatus on a heated platform (34–36 °C) for 10–15 min per day over 3 consecutive days prior to data collection. On the day of measurement, mice were placed in acrylic restrainers on the warmed platform to promote tail vasodilation and reduce motion artifacts. Each session consisted of 10 initial conditioning cycles followed by 30 measurement cycles, and values with movement artifacts or abnormal pressure curves were automatically excluded by the software. For each mouse, systolic, diastolic, and mean arterial pressures, along with heart rate, were calculated as the average of at least 10 valid measurement cycles. All recordings were performed at the same time of day to control for circadian variability. Data are reported as the mean of 2–3 independent sessions per animal.

### Behavioral Assays

For the Open Field Test, mice were placed in the center of a square arena and allowed to freely explore for 10 minutes, with behavior recorded by an overhead camera and analyzed using Noldus EthoVision XT software. Locomotor activity was quantified as total distance traveled, while anxiety-like behavior was measured as the percentage of time spent along the arena perimeter versus the center, with additional parameters including velocity, movement frequency, and immobility extracted for comprehensive assessment. The same arena was used for the Novel Object Recognition Test, which followed a four-day protocol: during habituation, mice were placed in the empty arena for 10 min on two consecutive days; on day three, two identical objects were positioned equidistant within the arena, and mice were allowed to explore for 20 minutes while movement trajectories and object interactions—defined as nose entries into a 2 cm zone around each object—were recorded with EthoVision. On day four, one of the familiar objects was replaced with a novel object of similar size but different shape and texture, and exploration was tracked for 20 minutes, with recognition memory quantified as a recognition index (time spent with the novel object relative to total object exploration). Standard exclusion criteria were applied, including removal of mice showing illness or injury, trials with tracking errors, sessions with <5% total exploration time or >80% immobility, or cases where both objects were not investigated. Arenas and objects were cleaned with ethanol between trials to minimize olfactory cues.

### Quantitative Polymerase Chain Reactions

Total RNA was isolated from cortical tissue using TRIzol reagent (Invitrogen) following the manufacturer’s protocol. Samples were placed on ice and mechanically homogenized in 1 ml of TRIzol. RNA concentration was determined using a BioTek Synergy LX Multimode Reader (Agilent). One microgram of RNA was reverse transcribed into cDNA using the High-Capacity cDNA Reverse Transcription Kit (Applied Biosystems). Quantitative PCR was performed with PowerUp SYBR Green Master Mix (Applied Biosystems) on a QuantStudio 3 Real-Time PCR System (Applied Biosystems). Data were analyzed using the ΔΔCt method, normalized to β2m, and expressed as fold change relative to wild-type naïve mice. Primers for semiquantitative and real-time PCR were obtained from Integrated DNA Technologies.

### Immunofluorescence of Human Brains

Use of postmortem human brain tissue was reviewed and deemed exempt by the Committee for the Protection of Human Subjects at the University of Texas Health Science Center at Houston. Written informed consent was obtained from the next of kin for all donors, in compliance with ethical guidelines and respect for patient autonomy. Cases included individuals with a clinical history of HFpEF and no evidence of co-existing neuropathologies such as Lewy body disease, TDP-43 pathology, glioblastoma, or other neurodegenerative conditions (e.g., Alzheimer’s or Huntington’s disease). Control tissue was obtained from individuals who died of non-cardiovascular, non-neurodegenerative causes. Formalin-fixed, paraffin-embedded blocks from the cingulate gyrus, a region known to undergo atrophy in HFpEF (9), were processed for histology. Sections were deparaffinized, rehydrated, and subjected to antigen retrieval by heating in citrate buffer (pH 6.0) for 30 minutes at 90 °C. After cooling, sections were blocked for 1 hour at room temperature in TBS containing 5% normal goat serum and 0.3% Triton X-100. Primary antibodies against IBA1 (1:500) and HIF-1α (1:100) were applied in blocking buffer and incubated overnight at 4 °C in a humidified chamber (IHC World). Following extensive washes in TBS with Tween, sections were incubated for 1 hour at room temperature in the dark with Alexa Fluor 594 donkey anti-rat and Alexa Fluor 647 donkey anti-rabbit secondary antibodies (all 1:500 in blocking buffer). Slides were then washed, treated with 1× TrueBlack in 70% ethanol for 1 minute to reduce autofluorescence, rinsed in TBS, and coverslipped with VECTASHIELD antifade mounting medium containing DAPI. Imaging was carried out at 20X or 63X magnification using a Leica DMi8 SPE confocal microscope. Representative images were adjusted uniformly for brightness and contrast with QuPath. Analyses were based on at least 15 fields across three sections per patient, with >100 cells evaluated per case.

### Immunofluorescence of Mouse Brains

At specified time points, mice were euthanized and perfused with PBS to clear circulating cells. Brains were removed and drop fixed in 4% PFA overnight at 4 °C, Following fixation, brains were rinsed in PBS for 2 hours and subsequently cryoprotected in PBS with 30% sucrose at 4 °C for 48–72 hours. Brains were embedded in Tissue-Tek O.C.T. compound (Sakura) and cryosectioned at 10 μm thickness using a Leica CM1860 UV cryostat. Sections were allowed to air dry at room temperature for 20 minutes, followed by PBS washing. For antigen retrieval, slides were incubated in citrate buffer (Sigma) for 30 min in a 90 °C water bath, then cooled to room temperature. Autofluorescence was quenched with 10 mM glycine containing 0.2% Triton X-100 in PBS for 1 hour at room temperature. Blocking was performed in PBS containing 5% BSA, 1% donkey serum, and 0.2% Triton X-100 for 1 h. Sections were then incubated overnight at 4 °C in a humidified chamber (IHC World) with primary antibodies against ASPA (1:300) or Iba1 (1:500) diluted in blocking buffer. After thorough PBS washes, secondary antibodies (1:500 in blocking buffer) were applied for 1 hour at room temperature in the dark. Slides were rinsed again in PBS and mounted using VECTASHIELD antifade medium with DAPI. Images were acquired at 20X or 63X magnification on a Leica DMi8 SPE confocal microscope. Representative images were uniformly adjusted for brightness and contrast, and cell quantification was performed with QuPath software. Analyses were based on at least three sections per sample, with more than 100 cells evaluated per condition.

### Statistical Analyses

Statistical analyses were conducted using GraphPad Prism 10 (GraphPad Software). Two-group comparisons were assessed with a two-tailed, unpaired t-test assuming a 95% confidence interval. For comparisons involving more than two groups or variables, one-way or two-way ANOVA was applied, also at a 95% confidence level, with Tukey’s post hoc test used for multiple comparisons where appropriate. Sample sizes for in vivo studies are noted in the figures and legends, reflecting pooled data from at least two independent experiments. For in vitro studies, sample sizes are similarly indicated and represent data from two or more independent replicates. Data are reported as Mean ± S.E.M. Statistical significance is defined in figure legends as \**P < 0*.*05*, \*\**P < 0*.*01*, \*\*\**P < 0*.*001*; results marked ‘ns’ denote non-significant findings.

## Acknowledgments

This work was supported by funding from the American Heart Association (https://doi.org/10.58275/AHA.25TPA1463689.pc.gr.233971 to MD), and McGovern Medical School Pilot Grant Program (to MD). These studies used the Small Animal Cardiovascular Phenotyping Service Center at UTHealth-Houston, sequencing services provided by Cancer Genomics Center at UTHealth-Houston (supported by CPRIT RP240610), and metabolomics services provided by the Metabolomics Facility at MD Anderson Cancer Center (supported in part by the University of Texas MD Anderson Cancer Center and P30CA016672).

